# Magneto-Photonic Gene Circuit for Minimally Invasive Control of Gene Expression in Mammalian Cells

**DOI:** 10.1101/2025.11.21.688514

**Authors:** E. Alejandro Castellanos Franco, Ryan Radawiec, Ashley Slaviero, Connor J. Grady, Brianna Ricker, Galit Pelled, Ute Hochgeschwender, Taeho Kim, Assaf A. Gilad

**Affiliations:** Department of Biomedical Engineering, Michigan State University, East Lansing, MI, United States; Department of Mechanical Engineering, Michigan State University, East Lansing, MI, United States; College of Medicine, Central Michigan University, Mount Pleasant, MI, United States; Department of Chemical Engineering and Materials Science, Michigan State University, East Lansing, MI, United States; Institute for Quantitative Health Science and Engineering, Michigan State University, East Lansing, MI, United States; Department of Radiology, Michigan State University, East Lansing, MI, United States; The Scojen Institute for Synthetic Biology, Reichman University, Herzliya, Israel

**Keywords:** Gene circuits, NanoLuc, Synthetic Biology, Optogenetics, Magnetogenetics, Electromagnetic perceptive gene (EPG)

## Abstract

Precise control of gene expression is one of the fundamental goals of synthetic biology. Whether the objective is to modify endogenous cellular function or induce the expression of molecules for diagnostic and therapeutic purposes, gene regulation remains a key aspect of biological systems. Over time, advances in protein engineering and molecular biology have led to the creation of gene circuits capable of inducing the expression of specific proteins in response to external stimulus such as light. These optogenetic, or light-activated circuits hold significant potential for gene therapy as a tool for regulating the expression of therapeutic genes within cells. However, the applications of optogenetic systems can be limited by the lack of efficient ways for light delivery inside cells or tissue. Our approach to address this challenge is to harness the power of bioluminescence to produce light directly inside cells using a luminescent enzyme. Combined with a photosensitive transcription factor, we report the development of a fully genetically encoded optogenetic circuit for control of gene expression. Furthermore, we utilized a magneto sensitive protein to engineer a split protein version of this luminescent enzyme, where its reconstitution is driven by a 50mT magnetic stimulus. Thus, resulting in a first-of-its-kind gene circuit activated by a combination of light and magnetic stimulus. We expect this work to advance the implementation of light-controlled systems without the need of external light sources, as well as serve as a basis for the development of future magneto-sensitive tools.

## INTRODUCTION

Gene circuits are an emergent technology that provides a novel approach to treating disease by using and controlling the expression of therapeutic genes^1–3^. Among these circuits, optogenetic systems are particularly interesting due to their ability to use light to modify cellular function with remarkable temporal and spatial resolution^4,5^. This is achieved by introducing naturally occurring or engineered light-sensitive proteins into cells that can either interact with endogenous pathways^2,5^, or act in tandem with other proteins to create brand new light-activated circuits^6^. One of such light-sensitive molecules is the bacterial photoreceptor EL222^7,8^. Originally found in *Erythrobacter litoralis*, EL222 is a light-sensing transcription factor containing a photosensing light-oxygen-voltage (LOV) domain and a DNA-binding helix-turn-helix (HTH) domain^8^. Exposure to light causes a conformational change within EL222 that allows it to dimerize and bind DNA which results in transcription of downstream genes^9^. While EL222 enables optogenetic control of gene expression via light, these systems often require external light sources to deliver the necessary light, which poses a significant challenge for implementation, particularly for *in vivo* settings. As such, the development of readily available and minimally invasive tools for light delivery inside tissue remains of great importance for future implementations of light-actuated systems^5^.

To address this challenge, we are harnessing the power of bioluminescence along with a magneto sensitive protein to engineer genetically encoded tools for light delivery to be used in an optogenetic circuit. Bioluminescence is regarded as a promising alternative light source in gene circuits by utilizing enzymes to produce light directly inside targeted cells^10,11^. Luminescent enzymes (luciferases) catalyze the oxidation of specific substrates and emit light, which in turn can activate light-sensitive proteins^10,12^. In recent years, much research has been dedicated to engineering luciferases and their substrates to achieve different kinetics, brighter emissions, or to shift emission wavelengths. Engineered luciferases have been widely used in imaging applications^13^ acting as gene expression reporters^10,11^ as well as part of biosensors to detect metabolic changes^14–16^.

Many luciferases have also been used in split protein systems^17^, whereby enzymes that are split into non-functional fragments reassemble into a functional unit when in proximity to one another^18–20^. One glutamate biosensor utilizes a split version of NanoLuc luciferase to report metabolic changes in glutamate concentration through dynamic changes in light emission^16^; these result from glutamate binding to a sensing domain, which drives the reconstitution of the split luciferase. Notably, as split protein technology evolves, new platforms to drive the reconstitution of split proteins using an external stimulus have emerged. One such is the Electromagnetic Perceptive Gene (EPG)^21–23^ protein, a novel gene found in the glass catfish, *Kryptopterus vitreolus*, a fish known to respond to magnetic fields^24,25^. In a recent study, EPG was successfully utilized to drive the reconstitution of split proteins in response to magnetic stimulation, as demonstrated in a split horseradish peroxidase and a split herpes simplex virus thymidine kinase^26^.

Here, we created a novel gene circuit based on a split-NanoLuc fusion protein, in which the reconstitution of the split enzyme is driven by a specific magnetic stimulus. This circuit, that serves as a “biological switch,” comprises three elements: a light source, a light-sensitive transcription factor, and a reporter gene. Thus, these genetically encoded tools enable minimally invasive activation of a genetic circuit for control of gene expression in tissue.

## RESULTS

### Using an LED array to characterize EL222-mediated transcription in mammalian cells

For the first iteration of our EL222 gene circuit, we utilized a previously developed adaptation of the EL222 photoreceptor to make it compatible with mammalian systems^7^. This modified photoreceptor incorporates a transcription activation domain (TAD) fused to the N-terminus of EL222 which allows it to induce gene expression in mammalian cells when exposed to blue light (450 nm). Along with EL222, we prepared a secreted alkaline phosphatase (SEAP) reporter inducible by binding of active EL222 to a 5xC120 region, the natural binding partner of EL222 (**Fig. 1A**). Initial testing in HEK293FT cells showed that, 24 hours after a period of light stimulation, cells expressing EL222 presented significantly higher amounts of SEAP reporter compared to those without EL222 (**Fig.1B**).

**Figure 1:**
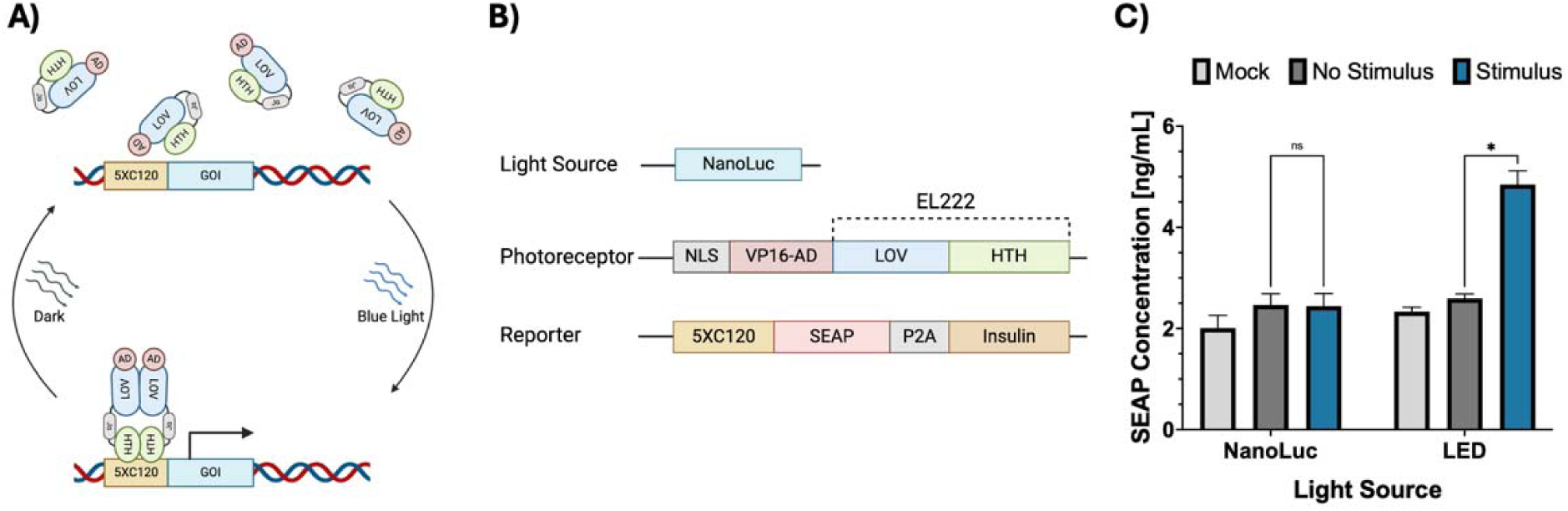
Initial assembly and testing of the EL222 circuit in HEK923FT cells. **(A)** Schematic of the EL222 transcription mechanism, blue light excitation (450 nm) causes a conformational change, dimerization, and DNA binding that is reversible in the absence of light. Binding of active EL222 to its 5xC120 partner sequence induces transcription of a downstream gene. **(B)** Components and structure of the EL222 circuit. The first iteration of this circuit consisted of three elements: a light source (cytosolic NanoLuc), a photoreceptor (VP16-EL222) and a reporter gene (5xC120 SEAP-P2A-INS). **(C)** SEAP reporter expression induced by LED or cytoplasmic NanoLuc. (*) Represents a P value < 0.05, calculated using an unpaired Student’s t-test.

After demonstrating that the circuit is activated by an LED light, we tested if the same circuit could be activated instead by the luminescent enzyme NanoLuc. We tested the hypothesis that, under the right conditions, a bright luciferase could produce enough light at the correct wavelength (450 nm) to activate EL222. In order to test this, we constructed a circuit consisting of three parts: a light source (NanoLuc), a photoreceptor (VP16-EL22), and a reporter (5xC120 SEAP). However, testing of the circuit described in **Fig.1A** shows that EL222 was activated by LED light but not by cytoplasmic NanoLuc (**Fig. 1C**). A possible explanation for this behavior is that, due to the much lower light intensity of luciferases compared to LEDs, other factors such as the distance between the light source and the photoreceptor, their spatial orientation and the exposure time play a more important role than we initially thought in the activation of EL222.

Following the initial testing, we focused on both achieving a better understanding of how EL222 responds to different input light strengths, as well as seeking ways to increase its response to lower light intensities. Thus, we engineered two new EL222 variants by replacing the existing VP16 TAD with either VP64 or VPR (transcription activation domains were cloned from Addgene plasmid #63798). These constructs were not meant to directly alter the sensitivity of EL222 to light stimulus; but instead, we sought to tune the response to light by changing its ability to induce transcription after activation. We used these three photoreceptor variants (VP16, VP64, and VPR) to study the effects of input light intensity (**Fig. 2B-F**) and stimulation time (**Fig. 2G-I**) on the transcriptional power of our circuit. By using an LED array controlled by an Arduino board (**Fig. 2A**), cells expressing EL222 were exposed to increasing levels of blue light LED over a constant stimulation time. Our results show that reporter expression increases with input strength as well as stimulation time, reaching a plateau at combinations of high light intensity and exposure time (**Fig. 2B, C**). The main conclusion from these experiments is that light intensity is crucial for optimal circuit activation. Given that improving the brightness of a luciferase is challenging, minimizing the distance between the luciferase and our photoreceptor, and finding optimal substrate concentrations become essential to achieve bioluminescent activation of EL222. This suggests that NanoLuc should be either fused or localized to the cell nucleus to minimize its distance to EL222^27,28^.

**Figure 2:**
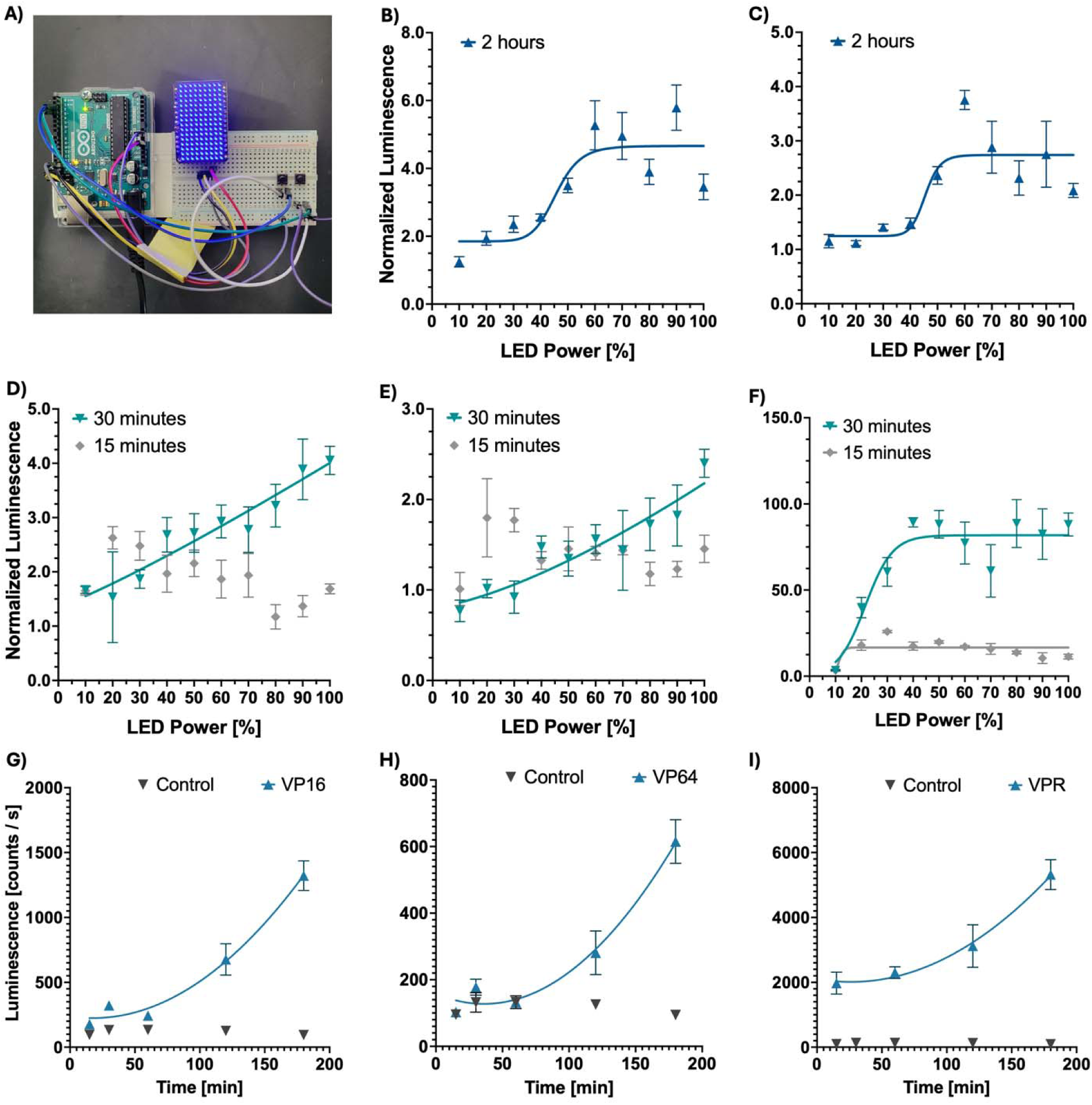
Characterization of EL222 constructs, light intensity and duration response. EL222 variants were obtained by replacing the VP16 transcription activation domain fused to EL222 with either VP64 or VPR activation domains. **(A)** Variable brightness LED matrix utilized for light stimulation and testing of EL222 variants. **(B-F)** Effect of light intensity on EL222-mediated transcription of the Firefly reporter gene. Cells expressing VP16-EL222 **(B, D)**, VP64-EL222 **(C, E)**, or VPR-EL222 **(F)** for 15 min, 30 min, or 2 h. Quantification of a Firefly reporter 24 hours post-stimulation showed levels of reporter expression that increase with LED strength. Regression analysis, represented by solid lines, shows the effect of LED strength on reporter expression can be approximated to a linear pattern at short exposure time or a sigmoidal pattern at longer time exposures. EL222-mediated Firefly expression at 60% LED intensity increased with stimulation time for cells expressing VP16-EL222 (**G**), VP64-EL222 (**H**), or VPR-EL222 (**I**).

### Developing a bioluminescent approach for EL222 activation using NanoLuc luciferase

Based on our findings, we developed two approaches to minimize the distance between EL222 and NanoLuc. In the first approach, we designed fusion constructs^27,29^ in which NanoLuc was fused to either the N-terminus or C-terminus of EL222 via flexible five-amino-acid-long linkers. Though this technique provided maximum proximity between the circuit elements, it also restricted the orientation and movement of its elements. Thus, for the second approach, we instead used an SV40 nuclear localization sequence to localize NanoLuc to the cell nucleus where EL222 is expressed (**Fig. 3B**). This method still provides a significant increase in proximity between circuit elements without compromising mobility.

**Figure 3:**
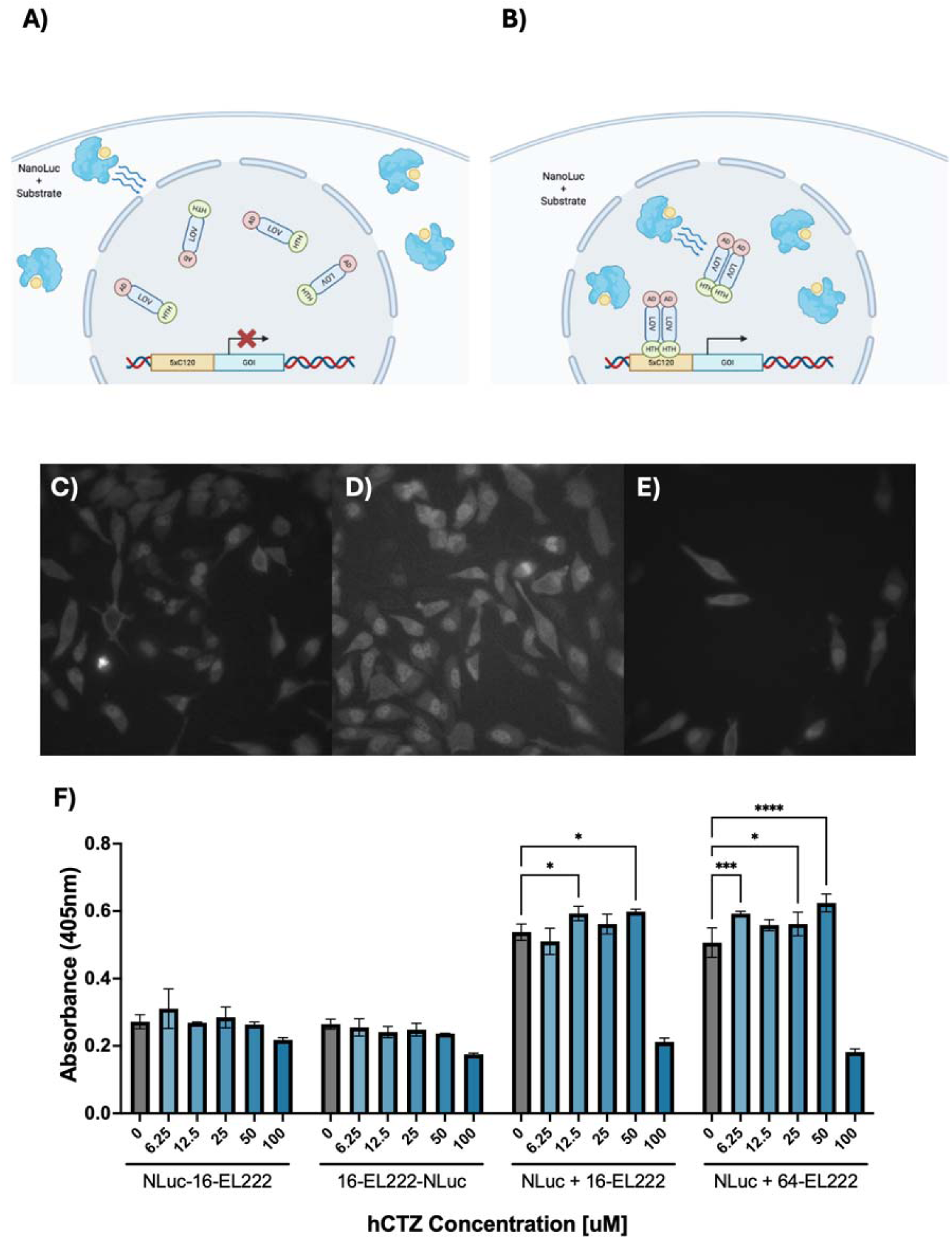
Bioluminescent approach for activation of EL222 in mammalian systems. **(A)** The distance between the cytoplasmic localization of the NanoLuc and the nuclear localization of EL222 implies that the distance hinders induced circuit activation. **(B)** Proposed an alternative approach to minimize the distance to EL222 by localizing the light source (NanoLuc) to the cell nucleus. **(C-E)** Bioluminescent imaging of several NanoLuc constructs used in this study. All constructs are nuclear-localized. **(C)** N-terminal NanoLuc-EL222 fusion, **(D)** C-terminal NanoLuc-EL222 fusion and **(E)** a nuclear localized NanoLuc variant. **(F)** SEAP reporter expression induced by different NanoLuc-EL222 fusion constructs or NanoLuc co-expressed with EL222 variants. Tested Cells were treated with hCTZ at concentrations ranging from 0 to 100 μM for 3 hours, followed by a media exchange. SEAP reporter activity was measured 12h post-stimulation. Reporter activity was quantified by measuring absorbance at 405 nm after a 72-h incubation with a pNPP SEAP substrate. SEAP reporter activity was detected in the groups transfected with nuclear-localized NanoLuc along with either VP16 or VP64-EL222. Data represents one out of three experiments; additional repetitions can be found in supplemental information (Fig S2). Statistical significance was calculated at a 5% significance level using Two-way analysis of variance (ANOVA) followed by Dunnett’s test. (*) = P < 0.05, (**) = P < 0.01, (***) = P < 0.001, (****) = P = < 0.0001.

We confirmed proper nuclear localization of our constructs using luminescence imaging (**Fig. 3C-E** and **Fig. S2**). Luminescent images were captured using a conventional inverted microscope equipped with an electron-multiplying CCD camera. To test these constructs, we designed new circuits utilizing NanoLuc-EL222 fusions, or a combination of nuclear-localized NanoLuc along with VP16-EL222 or VP64-EL222. We tested over a range of Coelenterazine-h (hCTZ) substrate concentrations and measured the resulting reporter production. Out of all tested constructs, VP64-EL222 showed the most consistent activation via NanoLuc illumination (**Fig. 3F**) across different experiments. These findings indicate that to activate the circuit using NanoLuc, a combination of proximity of the light source to the photoreceptor, a more potent transcriptional activator, and the appropriate substrate concentration is needed. Results also showed a decrease in reporter expression at 100μM hCTZ, which was correlated with a decrease in cell viability. Thus, we used this information to identify a suitable range of hCTZ concentrations to achieve EL222 activation without compromising cell survival.

### Characterization of the bioluminescence-activated optogenetic circuit

After confirming the activation of EL222 using luminescence, our focus moved to characterizing the existing circuit, beginning with optimizing the stimulation conditions. We previously learned that the intensity of the light, as well as the exposure time, are the key factors in determining the transcriptional activity of this circuit (**Fig. 2**). In the context of a luminescent enzyme, both of these factors are tied to the availability and concentration of the enzyme substrate. Therefore, we developed a series of assays in which we tested a circuit comprised of NLS-NanoLuc, VP64-EL222 and 5xC120 SEAP, over an array of substrate concentrations (0-50 µM hCTZ), number of substrate additions (1-3 additions) and stimulation duration (1.5h, 3h, 12h). Similar to the LED assays, these experiments showed that reporter expression increases with luciferase substrate concentration and stimulation time (**Fig. 4A**); which correlates to higher light intensity and longevity that results from higher availability of substrate. However, results also showed that higher concentrations of hCTZ resulted in overall lower reporter expression. This is likely the result of cell death, as cell viability assays (**Fig. 4B**) signal a decrease in cell survival at high hCTZ concentrations. Even though tuning stimulation conditions lead to a signal increase for the circuit, we sought to further enhance it by incorporating another variant of EL222, which utilizes the three-part TAD VPR instead of the previously used VP64. When stimulated using an LED, VPR shows drastically higher reporter expression compared to VP16 and VP64 (**Fig. 4C**), and when activated using our NanoLuc protocol, it shows significantly higher reporter expression than NanoLuc-activated VP64 (**Fig. 4D**).

**Figure 4:**
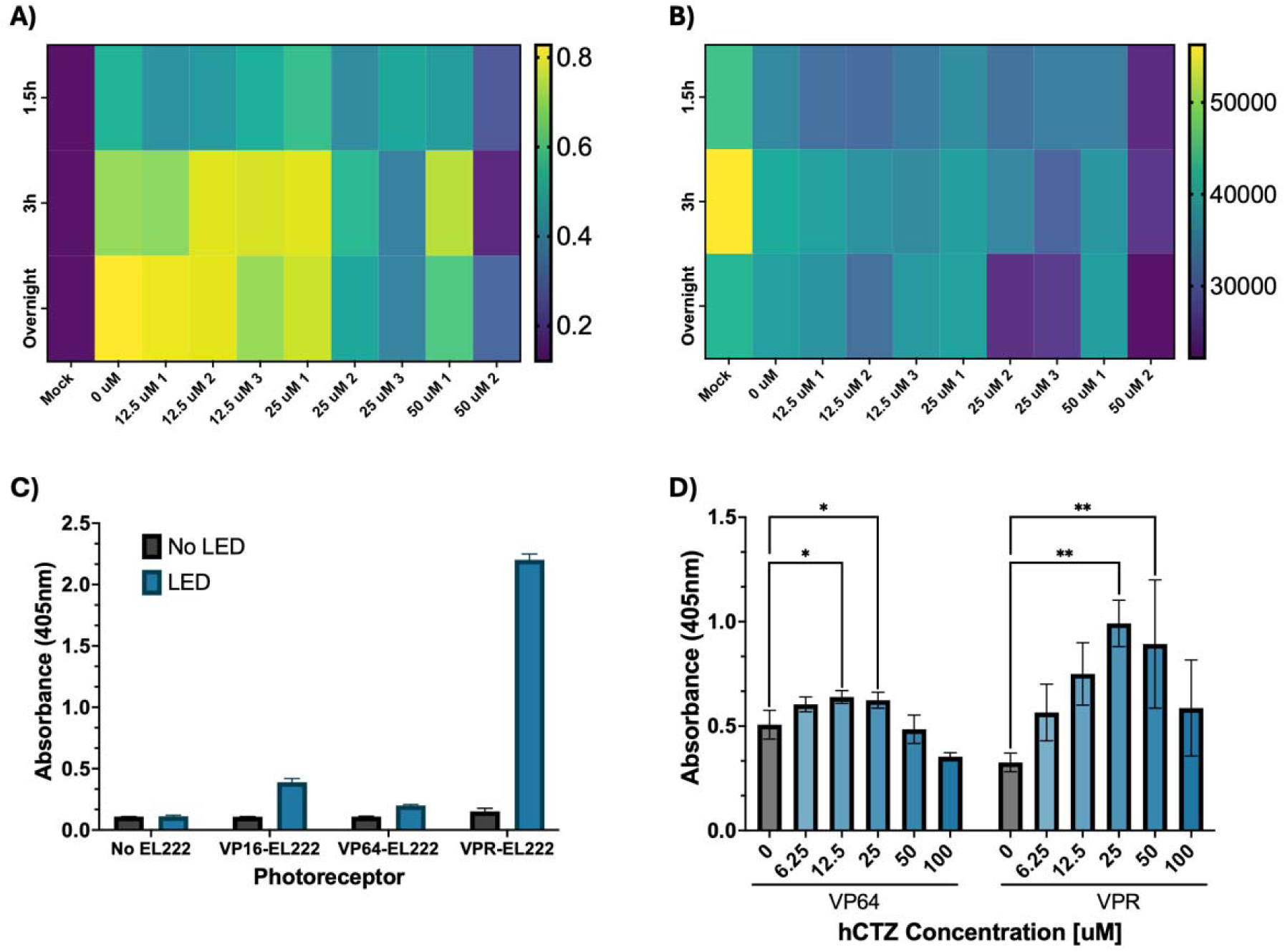
Optimizing Luminescent Activation of EL222 with NanoLuc. **(A)** Effect of substrate concentration, number of additions, and stimulation time on the EL222-mediated production of a SEAP reporter. Cells were provided with hCTZ ranging from 0-50uM, with some groups receiving subsequent additions of substrate in intervals of 30 min, up to a maximum of 3 additions, as denoted by the number following the concentration. The substrate was left for a period of 3 hours, and SEAP activity was measured the next morning. **(B)** Effect of substrate concentration, number of additions, and stimulation time on cell viability. Cell viability was determined via a cell titer blue assay; higher fluorescence denotes higher number of live cells. **(C)** Performance comparison of existing EL222 variants 24h post LED stimulation. **(D)** Performance of VP64-EL222 and VPR-EL222 using NanoLuc luciferase for activation over an array of hCTZ concentrations. Statistical significance was calculated at a 5% significance level using One-way analysis of variance (ANOVA) followed by Dunnett’s test. (*) = P < 0.05, (**) = P < 0.01, (***) = P < 0.001, (****) = P = < 0.0001.

### The electromagnetic perceptive protein controls gene expression via EL222

With the implementation of VPR-EL222, we developed a fully functional, genetically encoded circuit capable of controlling the expression of a gene of interest by adding luciferase substrate. At this stage, our goal became to incorporate another element that could act as an additional control for the circuit, reducing off-target activation by requiring two conditions to be met. To achieve this, we combined split-protein technology with an electromagnetic perceptive gene (EPG) protein to engineer a “biological switch” controlled by the presence of a magnetic stimulus.

This biological switch consists of two components: the first one is the EPG protein; a protein found in the glass catfish that responds to magnetic fields and that has been previously used as a platform for split protein reconstitution. The second element is NanoLuc luciferase, which we used extensively during circuit optimization and has several well-established split sites utilized in various applications. For this application, we utilized the “NanoBiT” split site as it allowed us to choose between fragments of different affinities for reconstitution. Thus, we designed a library of EPG-NanoLuc fusions constructs generally consisting of an EPG core flanked by the two fragments of split NanoLuc, fused to the N and C terminus of EPG by five amino acid-long linkers^26^. Each component had options to choose from, including two versions of EPG (full or truncated protein), two types of linkers (flexible or rigid), and three pairs of split NanoLuc fragments (native peptide, peptide 86 and peptide 114; see **Fig. S1** and **Table S1**).

The principle behind these constructs is to use EPG as a “hinge” to drive the reconstitution of split-NanoLuc, creating separate ON and OFF states that limit the amount of light produced, therefore acting as a second gate for the activation of EL222. We screened constructs by incorporating them into the EL222 circuit in place of full-length NanoLuc and comparing reporter expression levels between groups with or without a magnetic stimulus (**Fig. 5** and **Fig. S3**). Higher levels of reporter expression were associated with EPG-driven NanoLuc reconstitution. Through several rounds of screening, we identified eight main candidates for magneto reception in the initial library. During testing, we used an electromagnetic coil to deliver magnetic stimulus in the 50 mT range^22,23^ after addition of the luciferase substrate. From the constructs tested, only two constructs, RF114 and fRR114, showed a statistically significant difference in response to electromagnetic field (EMF).

**Figure 5:**
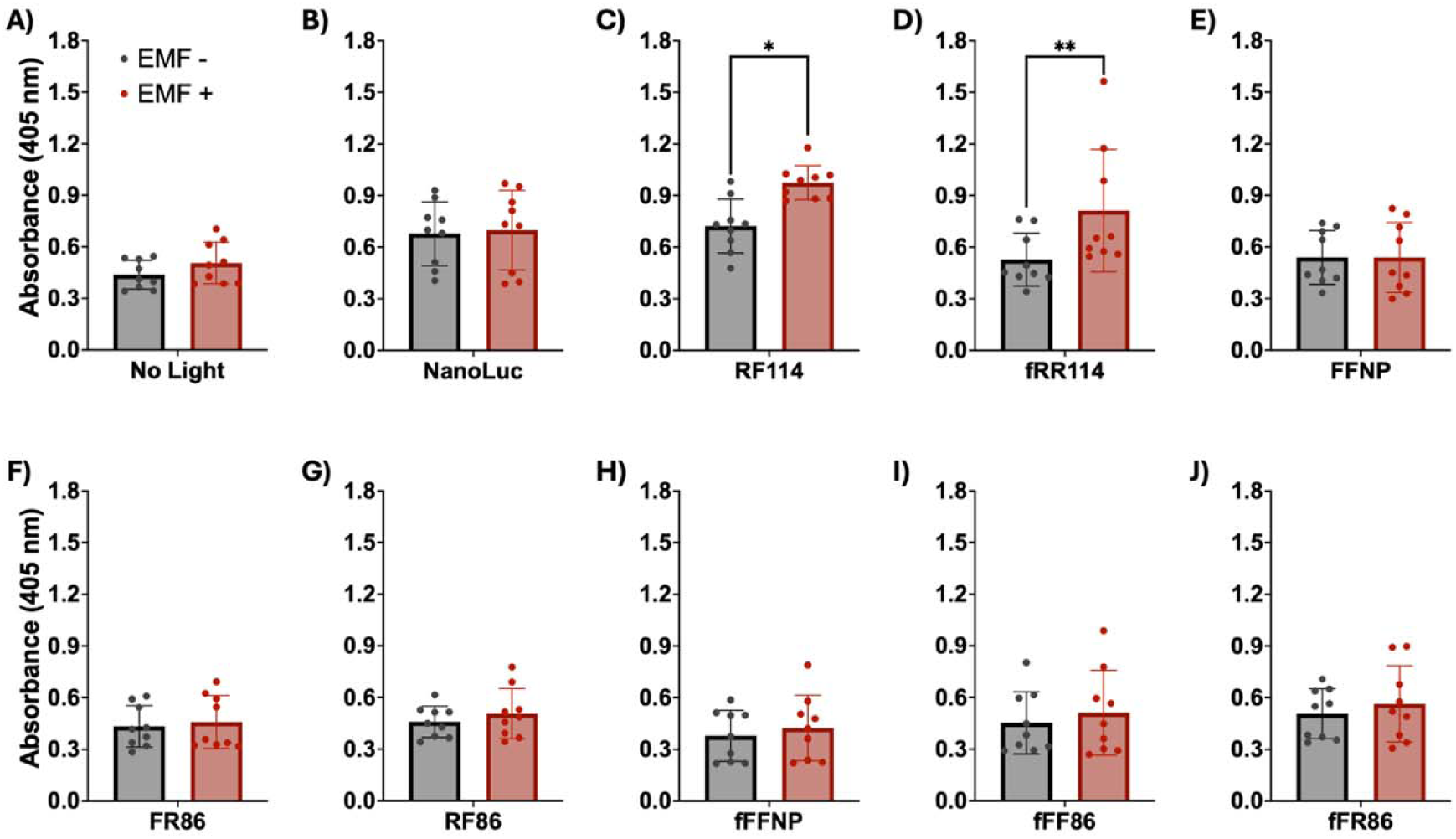
Control of the VPR-EL222 circuit using the magneto-receptive protein EPG. **(A-J)** Cells expressing one EPG-NanoLuc construct, VPR-EL222, and 5xC120 SEAP were treated with 25 μM hCTZ and placed in a dark incubator. One plate received three rounds of EMF pulses following a 15s ON 5min OFF pattern, each round separated by 2 hours. SEAP expression was measured 24 hours after stimulation. No light **(A)** group and NanoLuc **(B)** act as negative controls for EMF response. RF114 **(C)** and fRR114 **(D)** showed a significant increase in SEAP production following magnetic stimulus. Results shown represent an average of three independent experiments; each separate experiment contains information collected from three individual wells. Statistical significance was calculated at a 5% significance level using Two-way analysis of variance (ANOVA) followed by Sidak’s test. (*) = P < 0.05, (**) = P < 0.01, (***) = P < 0.001, (****) = P < 0.0001

The two EPG-driven NanoLuc constructs (RF114 and fRR114) that could activate the circuit and showed response to EMF were both based on the 114 variant of the SmBiT fragment, a variant known for its comparatively lower reconstitution affinity. In variant RF114, a truncated version of EPG (obtained by removing the translocation and GPI anchor signal sequences from EPG) was fused at the N-terminus to the LgBiT of NanoLuc using a rigid linker, and at the C-terminus to the SmBiT 114 using a flexible linker. This variant shows a 35.00% increase in absorbance compared to the control group associated with higher SEAP expression (P value = 0.0330, two-way ANOVA with Sidak correction). Similarly, construct fRR114 shows 53.87% increase in absorbance which was found to be significant (P value = 0.0099, two-way ANOVA with Sidak correction). Variant fRR114 incorporates a full-length version of EPG along with rigid linkers fusing both the LgBiT and SmBiT to the N and C-terminus, respectively. These findings indicate that the engineered EPG-NanoLuc requires both EMF and luciferase substrate for activation.

In summary, this study demonstrates the engineering of the first semi-synthetic magneto-photonic circuit for minimally invasive control of gene expression. This was done by combining split luciferase and a magnetosensitive protein to create a “bio-switch” that can produce light inside cells in the presence of its substrate and EMF. When coupled with a modified bacterial photoreceptor, this system can induce the expression of a desired gene in response to light.

## DISCUSSION

Here we present the first genetically encoded gene circuit that utilizes bioluminescence and electromagnetism to control the expression of a gene of interest. Our circuit draws upon previous work to adapt the bacterial photoreceptor EL222 to work in mammalian systems and expands this concept further by incorporating locally expressed NanoLuc luciferase to deliver light, overcoming the need for external light sources for activation. Moreover, we combined an electromagnetic preceptive gene (EPG) protein with NanoLuc to engineer a split luciferase whose reconstitution is driven by EPG in response to magnetic stimuli, acting as a “biological switch” for the circuit. This work represents a significant step forward in the implementation of optogenetic circuits, as it proposes a novel bioluminescent approach for their activation that overcomes common challenges while providing enhanced control and stability.

One of the core challenges of optogenetic systems lies in how to deliver the necessary amount of light where needed; this is particularly difficult inside tissue, where poor light penetration at certain wavelengths renders external light sources ineffective. We showed that, with proper calibration, this obstacle can be overcome using bioluminescence to produce light directly inside cells. Minimizing the distance between luciferase and photoreceptor was of utmost importance for this objective to compensate for the relatively lower light emission of luciferases compared to external light sources. These findings are in accordance with Adir et al, where they sought to address this issue by utilizing Bioluminescence Resonance Energy Transfer (BRET) to facilitate the activation of EL222 by Gaussia Luciferase in synthetic cells^28^. We anticipate that similar approaches can be used to activate other photoreceptors and that, as was shown with the inclusion of VPR, there is ample room for growth for this technology. We expect that the development of new and brighter luciferases, substrates, stronger activation domains and more sensitive photoreceptors will improve circuits like ours.

Gene circuits possess great therapeutic potential due to their ability to express therapeutic genes and molecules where they are required. However, these can also pose the risk of interfering with existing metabolic pathways, due to incorporating endogenous enzymes or metabolites as part of the circuit. This is a major challenge that we have addressed in our work by designing a synthetic gene circuit composed almost exclusively of non-mammalian proteins. By utilizing marine luciferases, fish genes, bacterial photoreceptors and viral activation domains, we greatly minimized the potential for crosstalk between our circuit and existing signaling pathways, therefore reducing harmful effects to the host cells. Furthermore, the use of light and magnetic fields as inputs also contributes to minimizing crosstalk, as neither of these elements is commonly involved in mammalian signaling pathways. The dual input system also presents the added benefit of reducing off-target activation of the circuit by requiring both light (produced by the addition of hCTZ) and EMF (necessary for split enzyme reconstitution) to be present for activation. Although all our work was done using in-vitro models, these properties that facilitate control and stability of the circuit are especially relevant for potential in vivo applications.

In conclusion, our main goal was to develop the first synthetic gene circuit controlled by bioluminescence and magnetism. We achieved this by designing a circuit based around the EL222 photoreceptor where the necessary light for activation was produced locally by a NanoLuc luciferase. Moreover, we engineered a split version of this enzyme fused to a magneto receptive protein (EPG) that has been previously used to drive the reconstitution of split proteins. The result was a “biological switch” that when provided with the substrate hCTZ and EMF stimulation, was able to activate EL222 and induce gene transcription. We expect that this approach will be useful for the implementation of other light-sensitive gene circuits, as well set up a framework for the utilization of magneto receptive proteins as regulatory elements in synthetic protein constructs and gene circuits.

## MATERIALS AND METHODS

### Cell culture conditions

LED stimulation experiments were done in HEK293FT or HeLa cells. HEK293FT cells were cultured in Dulbecco’s Modified Eagle Medium (DMEM) supplemented with 10% Fetal Bovine Serum (FBS), Penicillin-Streptomycin antibiotic and Geneticin G418 antibiotic. These cells were kept inside an incubator under 5% CO2 at 37C. HeLa cells were cultured under identical conditions using DMEM supplemented with 10% FBS and Penicillin-Streptomycin antibiotic. For LED experiments, HEK or HeLa cells were plated in white-wall clear-bottom 96-well plates at a density of 10,000 cells per well; and left in the incubator overnight prior to transfection.

### Split construct design and cloning

Split NanoLuc-EPG fusion constructs were designed based on the NanoBiT split luciferase system (Promega). NanoLuc was divided into Large BiT and Small BiT fragments which were fused to the N and C terminus of EPG respectively via five amino acid linkers. Additionally, an SV40 nuclear localization sequence was added to the N-terminus of the construct to ensure expression at the cell nucleus. To create a library of candidates for testing, each construct could incorporate different versions of its components; these include two versions of EPG, two kinds of linkers and three variants of the Small BiT. The two versions of EPG correspond to either a full EPG protein or a truncated version of EPG obtained by removing signaling sequences at both ends of the protein. Likewise, two types of linkers were used, either flexible (GGGGS) or rigid (PAPAP)^30^. Finally, three variants of the small peptide with varying affinities for its Large BiT partner were utilized: peptide 86 (high affinity), native peptide (neutral affinity), and peptide 114 (low affinity). 24 different constructs were cloned following this pattern via Gibson Assembly and introduced into pcDNA3.1(+) vectors for mammalian expression. Each fusion protein was named after its components, e.g. the name fRF86 denotes the use of full-EPG, a rigid linker connecting the large fragment, a flexible linker connecting the small fragment, and a peptide 86 small BiT.

### Transfection conditions

To facilitate light-induced transcription in mammalian cells, the photoreceptor and reporter elements of the EL222 circuit were expressed in the respective cell line via transient transfection. Transient transfections were done using a commercially available Lipofectamine 3000 kit; cells received a total of 100ng of DNA split between photoreceptor and reporter in a 1:1 ratio. The reporter of choice for this assay was a Firefly luciferase controlled by an inducible 5xC120 promoter. Three versions of the EL222 photoreceptor were tested during this experiment, each corresponding to EL222 fusions with different transcription activator (AD) domains: VP16, VP64, and VPR.

### Fabrication of a 9x16 LED Matrix with variable Brightness control

A 9x16 blue LED matrix was controlled using an Arduino Uno. The matrix was constructed with an Adafruit IS31FL3731 PWM LED driver breakout board populated with a 9x16 grid of blue LEDs. Power was supplied to the driver board directly from the Arduino Uno’s 3.3V and GND pins.

Manual adjustments of LED brightness could be made using two pushbuttons that integrated into the circuit. The two buttons were connected to digital pins D2 and D3 on the Arduino for brightness control. The Arduino communicated with the driver board using the I2C protocol, using pins A4 (SDA) and A5 (SCL), and was programmed using the Arduino IDE with the Adafruit_IS31FL3731 (v1.2.3), Adafruit_GFX, and Wire libraries. A custom script was developed to translate button presses, where one button increased the overall brightness of the LED matrix by approximately 10% (a value of 25 out of 255) and the other decreased it by the same amount. This functionality was achieved by adjusting the PWM value sent to all 144 LEDs through the driver.

### LED stimulation protocol

Two main protocols were developed to test the effect of light illumination on the EL222-mediated transcription in mammalian cells. To test the effect of input strength in reporter expression, cells expressing EL222 were placed inside a dark incubator (24h post-transfection) and the previously described LED array setup was used to deliver constant light stimulation for a set amount of time, over a range of LED power settings. Light intensity was modulated by regulating the power provided to the LED; a total of ten power settings were tested, ranging from 10% power (minimum light intensity) to 100% (max light intensity) in intervals of 10%. During a single experiment, all ten power settings were tested over a set stimulation time: 15min and 30 min (VPR construct), 15min 30min and 2 hours (VP16 and VP64 constructs).

Conversely, to test the time-dependence of the circuit, cells expressing EL222 were exposed to LED light at a constant intensity for variable amounts of time. Time-dependence experiments were done at 60% LED strength, and reporter expression was measured at five lengths of light stimulation: 15min, 30min, 1h, 2h, and 3h.

### Reporter quantification

Activation and performance of the EL222 circuit were determined by measuring the expression of a Firefly luciferase reporter. 24h post LED stimulation, cells that received light stimulation were incubated with D-luciferin for 15 min at 150mg/mL concentration; after incubation, luminescence was measured using a Tecan Spark plate reader. To avoid any crosstalk or background resulting from overlapping wavelengths between NanoLuc and Firefly, a Secreted Embryonic Alkaline Phosphatase (SEAP) reporter was chosen instead for assays where NanoLuc was used as a light source. The SEAP reporter was also modified to simultanously produce human insulin using a self-cleaving “P2A” peptide sequence. This provides an alternative for reporter detection, and possible future applications. SEAP quantification was done through a commercially available colorimetric assay (AnasSpec) 24h post-hCTZ addition. Samples were incubated with the corresponding substrate for 72h, after which absorbance at 405nm was measured using a Tecan Spark plate reader. Color change at the respective wavelength (clear to yellow) is proportional to the concentration of SEAP in the sample.

### Luminescence imaging

All luminescence imaging was done in HEK293FT cells transiently transfected with the appropriate NanoLuc/NanoLuc-EL222 plasmid and cultured on 35mm glass-bottom dishes. Prior to imaging, culture medium was replaced with Fluorobrite DMEM containing Fluorofurimazine (Promega) at 25μM final concentration. Imaging was done using an IX71 Inverted Microscope (Olympus) equipped with a UApo/340 40X Oil Iris objective lens (NA = 1.35) and an electron multiplying CCD (EM-CCD) iXON Ultra 888 camera (Oxford Technologies). Images were taken at 8s exposure time.

### Data processing

Data normalization of each experimental group was done using Microsoft Excel; data visualization, non-linear regression, and statistical analysis were performed using Prism 10 software. Statistical significance was calculated at a 5% significance level using either One-way or Two-way analysis of variance (ANOVA). (*) = P < 0.05, (**) = P < 0.01, (***) = P < 0.001, (****) = P = < 0.0001. Non-linear regression curves for the time-dependence experiment follow a second order polynomial, whereas input strength-dependence assays follow a four-parameter sigmoidal curve model.

## Author Contributions

E.A.C.F and A.A.G. designed experiments and wrote the manuscript. R.R., A.S., C.J.G., and B.R. provided assistance carrying out experiments. G.P., U.H., T.K., and A.A.G. supervised the project.

## ACKNOWLEDGEMENTS

Figures contain illustrations created with BioRender.com. A.A.G. acknowledges financial support from the NIH/NIBIB: R01-EB031008; R01-EB030565; R01-EB031936. We thank Dr. Kevin Gardner, CUNY, New York, NY, for providing plasmids pcDNA3.1-CAG-VP-EL222 and pcDNA3.1-5xC120-FireflyLuc.

