## Supplemental Figures for "Magneto-Photonic Gene Circuit for Minimally Invasive Control of Gene Expression in Mammalian Cells"

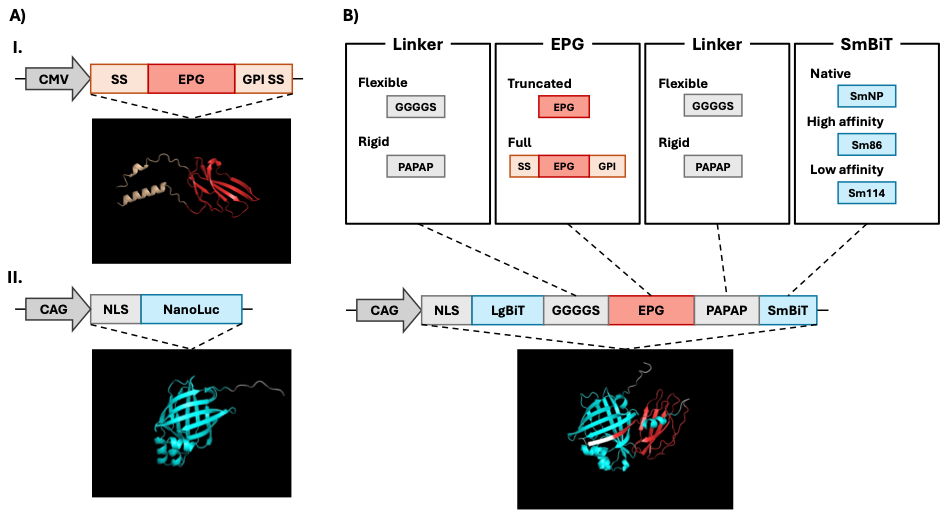

**Figure S1: Linear maps and structure prediction models for EPG (A-I), NanoLuc (A-II), and EPG-split-NanoLuc fusion constructs (B).** **(A-I)** The electromagnetic perceptive gene (EPG) protein shares a similar structure to members of the Ly6/uPAR family of proteins, characterized by their three-finger protein domain (red). EPG possesses an N-terminus translocation signal sequence and a C-terminus GPI anchor signal sequence. **(A-II)** NanoLuc luciferase is a luminescent enzyme possessing a b-barrel structure that catalyzes the oxidation of coelenterazine, or its derivatives, producing blue light (460nm) in the process. This version of NanoLuc was fused to an SV40 nuclear localization sequence to improve compatibility with the EL222 system. **(B)** General structure of EPG-NanoLuc fusion constructs. Synthetic fusions were built by fusing split-NanoLuc fragments (large subunit and small subunit) to EPG via 5-amino acid linkers. Each synthetic protein results from the combination of four variable factors: first linker flexibility (flexible or rigid), EPG variant (full or truncated), second linker flexibility (flexible or rigid) and small subunit affinity (native, low or high).

**Table S1: Components of EPG-NanoLuc fusion proteins cloned during this study.** A total of 24 distinct constructs were prepared and tested, accounting for every possible combination of the four variable factors. Each protein received a unique identifier that summarizes the specific combination of factors utilized (e.g. fFRNP: full EPG, flexible first linker, rigid second linker, and native variant small subunit).

| Construct | First Linker | Magnetoreceptor | Second Linker | SmBiT Variant |
| --- | --- | --- | --- | --- |
| FFNP | Flexible | Truncated EPG | Flexible | Native peptide |
| FRNP | Flexible | Truncated EPG | Rigid | Native peptide |
| RFNP | Rigid | Truncated EPG | Flexible | Native peptide |
| RRNP | Rigid | Truncated EPG | Rigid | Native peptide |
| FF86 | Flexible | Truncated EPG | Flexible | Peptide 86 |
| FR86 | Flexible | Truncated EPG | Rigid | Peptide 86 |
| RF86 | Rigid | Truncated EPG | Flexible | Peptide 86 |
| RR86 | Rigid | Truncated EPG | Rigid | Peptide 86 |
| FF114 | Flexible | Truncated EPG | Flexible | Peptide 114 |
| FR114 | Flexible | Truncated EPG | Rigid | Peptide 114 |
| RF114 | Rigid | Truncated EPG | Flexible | Peptide 114 |
| RR114 | Rigid | Truncated EPG | Rigid | Peptide 114 |
| fFFNP | Flexible | EPG | Flexible | Native peptide |
| fFRNP | Flexible | EPG | Rigid | Native peptide |
| fRFNP | Rigid | EPG | Flexible | Native peptide |
| fRRNP | Rigid | EPG | Rigid | Native peptide |
| fFF86 | Flexible | EPG | Flexible | Peptide 86 |
| fFR86 | Flexible | EPG | Rigid | Peptide 86 |
| fRF86 | Rigid | EPG | Flexible | Peptide 86 |
| fRR86 | Rigid | EPG | Rigid | Peptide 86 |
| fFF114 | Flexible | EPG | Flexible | Peptide 114 |
| fFR114 | Flexible | EPG | Rigid | Peptide 114 |
| fRF114 | Rigid | EPG | Flexible | Peptide 114 |
| fRR114 | Rigid | EPG | Rigid | Peptide 114 |

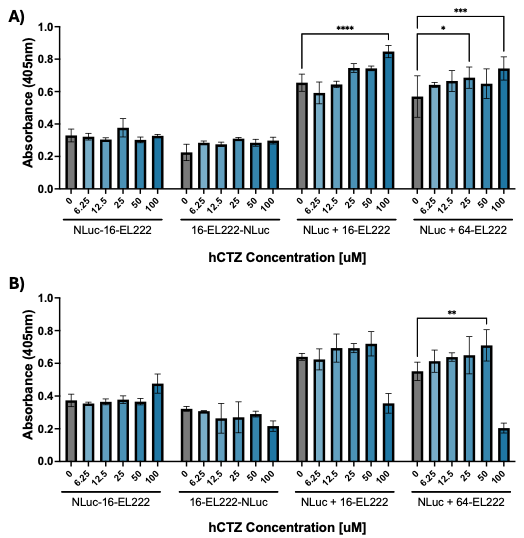

**Figure S2: Replicates of substrate concentration experiments for luminescent activation of EL222.** Quantification of SEAP reporter expression induced by NanoLuc-EL222 fusions or NanoLuc co-expressed with EL222 variants. Statistical significance was calculated at a 5% significance level using Two-way analysis of variance (ANOVA) followed by Dunnett’s test. (*) = P < 0.05, (**) = P < 0.01, (***) = P < 0.001, (****) = P = < 0.0001.

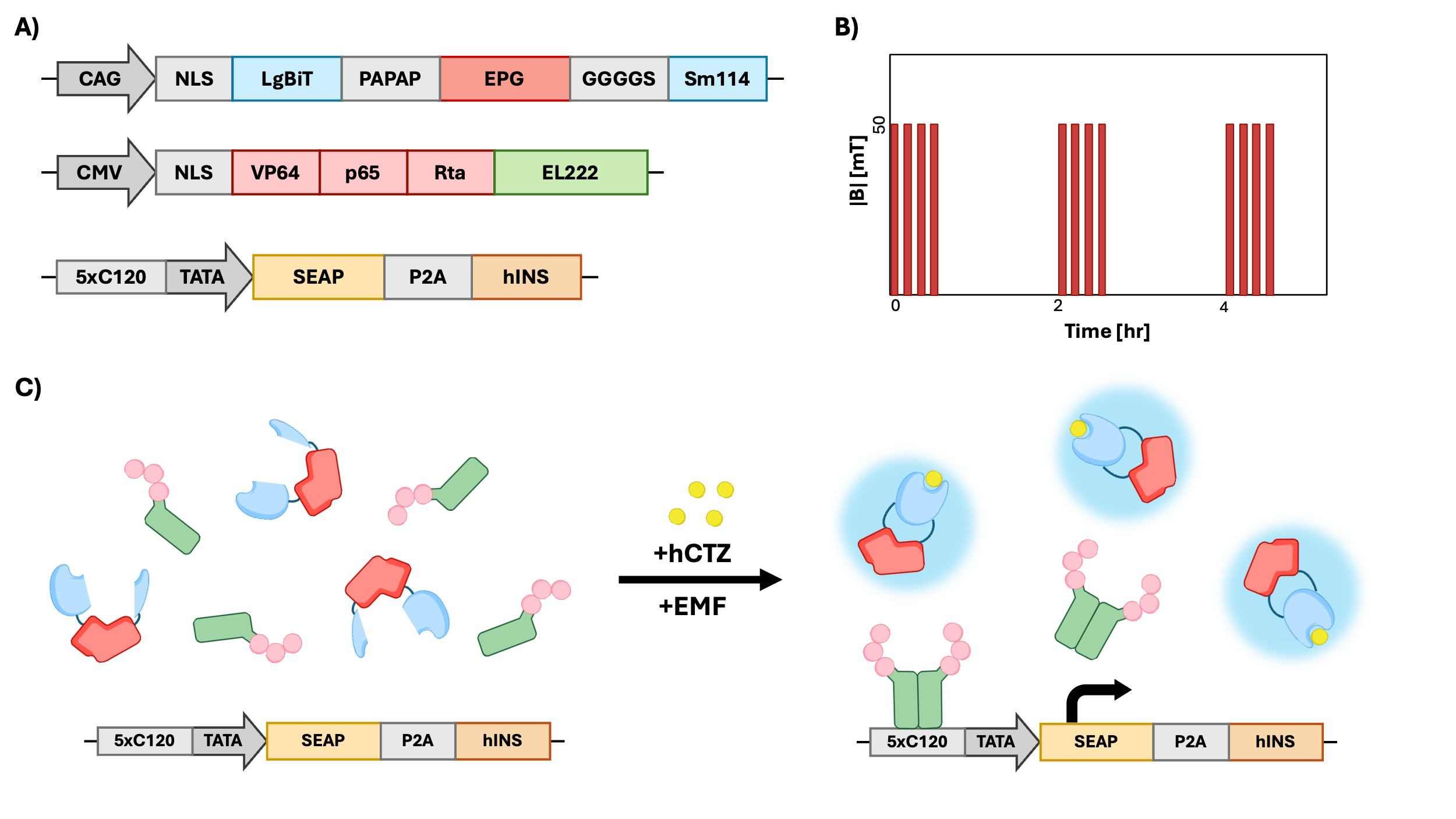

**Figure S3: Final design and elements of the magneto-photonic EL222 circuit.** **(A)** Linear map representations of the three circuit components. Nuclear-localized EPG-NanoLuc serves as both an intracellular light source and magneto sensitive component, VPR-EL222 functions as a photo-sensing component and transcription factor, and 5xC120 SEAP acts as inducible reporter for the system. **(B)** Visual representation of the electromagnetic stimulation pattern used in this study. After addition of hCTZ, cells received four magnetic stimulation pulses (50mT) following a 15s-ON 5min-OFF pattern; this is repeated two more times, each separated by a 2-hour interval. **(C)** Illustration of the magneto-photonic circuit during the inactive and active state. Magnetic stimulation induces EPG-driven reconstitution of NanoLuc, which produces blue light in the presence of hCTZ. As a results, EL222 dimerizes, bind DNA and promotes transcription of the reporter.

**Sequences of Constructs Used in This Study**

**VP16 EL222** – Vector: pcDNA3.1 – Promoter: CMV

Start-VP16-EL222-End

ATGGGCCCTAAAAAGAAGCGTAAAGTCGCCCCCCCGACCGATGTCAGCCTGGGGGACGAGCTCCACTTAGACGGCGAGGACGTGGCGATGGCGCATGCCGACGCGCTAGACGATTTCGATCTGGACATGTTGGGGGACGGGGATTCCCCGGGGCCGGGATTTACCCCCCACGACTCCGCCCCCTACGGCGCTCTGGATATGGCCGACTTCGAGTTTGAGCAGATGTTTACCGATGCCCTTGGAATTGACGAGTACGGTGGGGAATTCGGGGCAGACGACACACGCGTTGAGGTGCAACCGCCGGCGCAGTGGGTCCTCGACCTGATCGAGGCCAGCCCGATCGCATCGGTCGTGTCCGATCCGCGTCTCGCCGACAATCCGCTGATCGCCATCAACCAGGCCTTCACCGACCTGACCGGCTATTCCGAAGAAGAATGCGTCGGCCGCAATTGCCGATTCCTGGCAGGTTCCGGCACCGAGCCGTGGCTGACCGACAAGATCCGCCAAGGCGTGCGCGAGCACAAGCCGGTGCTGGTCGAGATCCTGAACTACAAGAAGGACGGCACGCCGTTCCGCAATGCCGTGCTCGTTGCACCGATCTACGATGACGACGACGAGCTTCTCTATTTCCTCGGCAGCCAGGTCGAAGTCGACGACGACCAGCCCAACATGGGCATGGCGCGCCGCGAACGCGCCGCGGAAATGCTCAGGACGCTGTCGCCGCGCCAGCTCGAGGTTACGACGCTGGTGGCATCGGGCTTGCGCAACAAGGAAGTGGCGGCCCGGCTCGGCCTGTCGGAGAAAACCGTCAAGATGCACCGCGGGCTGGTGATGGAAAAGCTCAACCTGAAGACCAGTGCCGATCTGGTGCGCATTGCCGTCGAAGCCGGAATCTAA

**VP64 EL222** – Vector: pcDNA3.1 – Promoter: CMV

Start-VP64-EL222-End

ATGGGCCCTAAAAAGAAGCGTAAAGTCGACGCATTGGACGATTTTGATCTGGATATGCTGGGAAGTGACGCCCTCGATGATTTTGACCTTGACATGCTTGGTTCGGATGCCCTTGATGACTTTGACCTCGACATGCTCGGCAGTGACGCCCTTGATGATTTCGACCTGGACATGCTGGAATTCGGGGCAGACGACACACGCGTTGAGGTGCAACCGCCGGCGCAGTGGGTCCTCGACCTGATCGAGGCCAGCCCGATCGCATCGGTCGTGTCCGATCCGCGTCTCGCCGACAATCCGCTGATCGCCATCAACCAGGCCTTCACCGACCTGACCGGCTATTCCGAAGAAGAATGCGTCGGCCGCAATTGCCGATTCCTGGCAGGTTCCGGCACCGAGCCGTGGCTGACCGACAAGATCCGCCAAGGCGTGCGCGAGCACAAGCCGGTGCTGGTCGAGATCCTGAACTACAAGAAGGACGGCACGCCGTTCCGCAATGCCGTGCTCGTTGCACCGATCTACGATGACGACGACGAGCTTCTCTATTTCCTCGGCAGCCAGGTCGAAGTCGACGACGACCAGCCCAACATGGGCATGGCGCGCCGCGAACGCGCCGCGGAAATGCTCAGGACGCTGTCGCCGCGCCAGCTCGAGGTTACGACGCTGGTGGCATCGGGCTTGCGCAACAAGGAAGTGGCGGCCCGGCTCGGCCTGTCGGAGAAAACCGTCAAGATGCACCGCGGGCTGGTGATGGAAAAGCTCAACCTGAAGACCAGTGCCGATCTGGTGCGCATTGCCGTCGAAGCCGGAATCTAA

**VPR EL222** – Vector: pcDNA3.1 – Promoter: CMV

Start-VPR-EL222-End

ATGGGCCCTAAAAAGAAGCGTAAAGTCGACGCATTGGACGATTTTGATCTGGATATGCTGGGAAGTGACGCCCTCGATGATTTTGACCTTGACATGCTTGGTTCGGATGCCCTTGATGACTTTGACCTCGACATGCTCGGCAGTGACGCCCTTGATGATTTCGACCTGGACATGCTGATTAACTCTAGAAGTTCCGGATCTCCGAAAAAGAAACGCAAAGTTGGTAGCCAGTACCTGCCCGACACCGACGACCGGCACCGGATCGAGGAAAAGCGGAAGCGGACCTACGAGACATTCAAGAGCATCATGAAGAAGTCCCCCTTCAGCGGCCCCACCGACCCTAGACCTCCACCTAGAAGAATCGCCGTGCCCAGCAGATCCAGCGCCAGCGTGCCAAAACCTGCCCCCCAGCCTTACCCCTTCACCAGCAGCCTGAGCACCATCAACTACGACGAGTTCCCTACCATGGTGTTCCCCAGCGGCCAGATCTCTCAGGCCTCTGCTCTGGCTCCAGCCCCTCCTCAGGTGCTGCCTCAGGCTCCTGCTCCTGCACCAGCTCCAGCCATGGTGTCTGCACTGGCTCAGGCACCAGCACCCGTGCCTGTGCTGGCTCCTGGACCTCCACAGGCTGTGGCTCCACCAGCCCCTAAACCTACACAGGCCGGCGAGGGCACACTGTCTGAAGCTCTGCTGCAGCTGCAGTTCGACGACGAGGATCTGGGAGCCCTGCTGGGAAACAGCACCGATCCTGCCGTGTTCACCGACCTGGCCAGCGTGGACAACAGCGAGTTCCAGCAGCTGCTGAACCAGGGCATCCCTGTGGCCCCTCACACCACCGAGCCCATGCTGATGGAATACCCCGAGGCCATCACCCGGCTCGTGACAGGCGCTCAGAGGCCTCCTGATCCAGCTCCTGCCCCTCTGGGAGCACCAGGCCTGCCTAATGGACTGCTGTCTGGCGACGAGGACTTCAGCTCTATCGCCGATATGGATTTCTCAGCCTTGCTGGGCTCTGGCAGCGGCAGCCGGGATTCCAGGGAAGGGATGTTTTTGCCGAAGCCTGAGGCCGGCTCCGCTATTAGTGACGTGTTTGAGGGCCGCGAGGTGTGCCAGCCAAAACGAATCCGGCCATTTCATCCTCCAGGAAGTCCATGGGCCAACCGCCCACTCCCCGCCAGCCTCGCACCAACACCAACCGGTCCAGTACATGAGCCAGTCGGGTCACTGACCCCGGCACCAGTCCCTCAGCCACTGGATCCAGCGCCCGCAGTGACTCCCGAGGCCAGTCACCTGTTGGAGGATCCCGATGAAGAGACGAGCCAGGCTGTCAAAGCCCTTCGGGAGATGGCCGATACTGTGATTCCCCAGAAGGAAGAGGCTGCAATCTGTGGCCAAATGGACCTTTCCCATCCGCCCCCAAGGGGCCATCTGGATGAGCTGACAACCACACTTGAGTCCATGACCGAGGATCTGAACCTGGACTCACCCCTGACCCCGGAATTGAACGAGATTCTGGATACCTTCCTGAACGACGAGTGCCTCTTGCATGCCATGCATATCAGCACAGGACTGTCCATCTTCGACACATCTCTGTTTGAATTCGGGGCAGACGACACACGCGTTGAGGTGCAACCGCCGGCGCAGTGGGTCCTCGACCTGATCGAGGCCAGCCCGATCGCATCGGTCGTGTCCGATCCGCGTCTCGCCGACAATCCGCTGATCGCCATCAACCAGGCCTTCACCGACCTGACCGGCTATTCCGAAGAAGAATGCGTCGGCCGCAATTGCCGATTCCTGGCAGGTTCCGGCACCGAGCCGTGGCTGACCGACAAGATCCGCCAAGGCGTGCGCGAGCACAAGCCGGTGCTGGTCGAGATCCTGAACTACAAGAAGGACGGCACGCCGTTCCGCAATGCCGTGCTCGTTGCACCGATCTACGATGACGACGACGAGCTTCTCTATTTCCTCGGCAGCCAGGTCGAAGTCGACGACGACCAGCCCAACATGGGCATGGCGCGCCGCGAACGCGCCGCGGAAATGCTCAGGACGCTGTCGCCGCGCCAGCTCGAGGTTACGACGCTGGTGGCATCGGGCTTGCGCAACAAGGAAGTGGCGGCCCGGCTCGGCCTGTCGGAGAAAACCGTCAAGATGCACCGCGGGCTGGTGATGGAAAAGCTCAACCTGAAGACCAGTGCCGATCTGGTGCGCATTGCCGTCGAAGCCGGAATCTAA

**5xC120 Firefly** – Vector: pcDNA3.1 – Promoter: 5xC120

Start-Firefly-End

ATGGAAGATGCCAAAAACATTAAGAAGGGCCCAGCGCCATTCTACCCACTCGAAGACGGGACCGCCGGCGAGCAGCTGCACAAAGCCATGAAGCGCTACGCCCTGGTGCCCGGCACCATCGCCTTTACCGACGCACATATCGAGGTGGACATTACCTACGCCGAGTACTTCGAGATGAGCGTTCGGCTGGCAGAAGCTATGAAGCGCTATGGGCTGAATACAAACCATCGGATCGTGGTGTGCAGCGAGAATAGCTTGCAGTTCTTCATGCCCGTGTTGGGTGCCCTGTTCATCGGTGTGGCTGTGGCCCCAGCTAACGACATCTACAACGAGCGCGAGCTGCTGAACAGCATGGGCATCAGCCAGCCCACCGTCGTATTCGTGAGCAAGAAAGGGCTGCAAAAGATCCTCAACGTGCAAAAGAAGCTACCGATCATACAAAAGATCATCATCATGGATAGCAAGACCGACTACCAGGGCTTCCAAAGCATGTACACCTTCGTGACTTCCCATTTGCCACCCGGCTTCAACGAGTACGACTTCGTGCCCGAGAGCTTCGACCGGGACAAAACCATCGCCCTGATCATGAACAGTAGTGGCAGTACCGGATTGCCCAAGGGCGTAGCCCTACCGCACCGCACCGCTTGTGTCCGATTCAGTCATGCCCGCGACCCCATCTTCGGCAACCAGATCATCCCCGACACCGCTATCCTCAGCGTGGTGCCATTTCACCACGGCTTCGGCATGTTCACCACGCTGGGCTACTTGATCTGCGGCTTTCGGGTCGTGCTCATGTACCGCTTCGAGGAGGAGCTATTCTTGCGCAGCTTGCAAGACTATAAGATTCAATCTGCCCTGCTGGTGCCCACACTATTTAGCTTCTTCGCTAAGAGCACTCTCATCGACAAGTACGACCTAAGCAACTTGCACGAGATCGCCAGCGGCGGGGCGCCGCTCAGCAAGGAGGTAGGTGAGGCCGTGGCCAAACGCTTCCACCTACCAGGCATCCGCCAGGGCTACGGCCTGACAGAAACAACCAGCGCCATTCTGATCACCCCCGAAGGGGACGACAAGCCTGGCGCAGTAGGCAAGGTGGTGCCCTTCTTCGAGGCTAAGGTGGTGGACTTGGACACCGGTAAGACACTGGGTGTGAACCAGCGCGGCGAGCTGTGCGTCCGTGGCCCCATGATCATGAGCGGCTACGTTAACAACCCCGAGGCTACAAACGCTCTCATCGACAAGGACGGCTGGCTGCACAGCGGCGACATCGCCTACTGGGACGAGGACGAGCACTTCTTCATCGTGGACCGGCTGAAGAGCCTGATCAAATACAAGGGCTACCAGGTAGCCCCAGCCGAACTGGAGAGCATCCTGCTGCAACACCCCAACATCTTCGACGCCGGGGTCGCCGGCCTGCCCGACGACGATGCCGGCGAGCTGCCCGCCGCAGTCGTCGTGCTGGAACACGGTAAAACCATGACCGAGAAGGAGATCGTGGACTATGTGGCCAGCCAGGTTACAACCGCCAAGAAGCTGCGCGGTGGTGTTGTGTTCGTGGACGAGGTGCCTAAAGGACTGACCGGCAAGTTGGACGCCCGCAAGATCCGCGAGATTCTCATTAAGGCCAAGAAGGGCGGCAAGATCGCCGTGTAA

**5xC120 SEAP-P2A-hINS** – Vector: pcDNA3.1 – Promoter: 5xC120

Start-SEAP-P2A-hINS-End

ATGCTGGGGCCCTGCATGCTGCTGCTGCTGCTGCTGCTGGGCCTGAGGCTACAGCTCTCCCTGGGCATCATCCCAGTTGAGGAGGAGAACCCGGACTTCTGGAACCGCGAGGCAGCCGAGGCCCTGGGTGCCGCCAAGAAGCTGCAGCCTGCACAGACAGCCGCCAAGAACCTCATCATCTTCCTGGGCGATGGGATGGGGGTGTCTACGGTGACAGCTGCCAGGATCCTAAAAGGGCAGAAGAAGGACAAACTGGGGCCTGAGATACCCCTGGCCATGGACCGCTTCCCATATGTGGCTCTGTCCAAGACATACAATGTAGACAAACATGTGCCAGACAGTGGAGCCACAGCCACGGCCTACCTGTGCGGGGTCAAGGGCAACTTCCAGACCATTGGCTTGAGTGCAGCCGCCCGCTTTAACCAGTGCAACACGACACGCGGCAACGAGGTCATCTCCGTGATGAATCGGGCCAAGAAAGCAGGGAAGTCAGTGGGAGTGGTAACCACCACACGAGTGCAGCACGCCTCGCCAGCCGGCACCTACGCCCACACGGTGAACCGCAACTGGTACTCGGACGCCGACGTGCCTGCCTCGGCCCGCCAGGAGGGGTGCCAGGACATCGCTACGCAGCTCATCTCCAACATGGACATTGACGTGATCCTAGGTGGAGGCCGAAAGTACATGTTTCGCATGGGAACCCCAGACCCTGAGTACCCAGATGACTACAGCCAAGGTGGGACCAGGCTGGACGGGAAGAATCTGGTGCAGGAATGGCTGGCGAAGCGCCAGGGTGCCCGGTATGTGTGGAACCGCACTGAGCTCATGCAGGCTTCCCTGGACCCGTCTGTGACCCATCTCATGGGTCTCTTTGAGCCTGGAGACATGAAATACGAGATCCACCGAGACTCCACACTGGACCCCTCCCTGATGGAGATGACAGAGGCTGCCCTGCGCCTGCTGAGCAGGAACCCCCGCGGCTTCTTCCTCTTCGTGGAGGGTGGTCGCATCGACCATGGTCATCATGAAAGCAGGGCTTACCGGGCACTGACTGAGACGATCATGTTCGACGACGCCATTGAGAGGGCGGGCCAGCTCACCAGCGAGGAGGACACGCTGAGCCTCGTCACTGCCGACCACTCCCACGTCTTCTCCTTCGGAGGCTACCCCCTGCGAGGGAGCTCCATCTTCGGGCTGGCCCCTGGCAAGGCCCGGGACAGGAAGGCCTACACGGTCCTCCTATACGGAAACGGTCCAGGCTATGTGCTCAAGGACGGCGCCCGGCCGGATGTTACCGAGAGCGAGAGCGGGAGCCCCGAGTATCGGCAGCAGTCAGCAGTGCCCCTGGACGAAGAGACCCACGCAGGCGAGGACGTGGCGGTGTTCGCGCGCGGCCCGCAGGCGCACCTGGTTCACGGCGTGCAGGAGCAGACCTTCATAGCGCACGTCATGGCCTTCGCCGCCTGCCTGGAGCCCTACACCGCCTGCGACCTGGCGCCCCCCGCCGGCACCACCGACGCCGCGCACCCGGGTCGTCGCAAGCGTGGAAGCGGAGCTACTAACTTCAGCCTGCTGAAGCAGGCTGGAGACGTGGAGGAGAACCCTGGACCTGTCGAATTCATGGCCCTCTGGATGAGGCTGCTTCCACTTCTTGCGCTCCTGGCGTTGTGGGGACCTGATCCGGCGGCAGCGTTCGTCAATCAGCATCTGTGCGGGAGTCACCTTGTCGAAGCATTGTACCTTGTTTGTGGAGAGCGCGGTTTTTTCTATACGCCTAAGACCCGCAGAGAGGCTGAAGATTTGCAAGTGGGACAAGTGGAGCTTGGAGGAGGGCCGGGTGCCGGTTCCCTCCAGCCTTTGGCTCTTGAGGGGTCCCTTCAGAAACGCGGGATAGTCGAACAATGTTGCACAAGCATATGCTCACTTTACCAACTGGAAAATTACTGCAACTAA

**NLS NanoLuc** – Vector: pcDNA3.1 – Promoter: CAG

Start-NLS-NanoLuc-End

ATGGGCCCTAAAAAGAAGCGTAAAGTCGTCTTCACACTCGAAGATTTCGTTGGGGACTGGCGACAGACAGCCGGCTACAACCTGGACCAAGTCCTTGAACAGGGAGGTGTGTCCAGTTTGTTTCAGAATCTCGGGGTGTCCGTAACTCCGATCCAAAGGATTGTCCTGAGCGGTGAAAATGGGCTGAAGATCGACATCCATGTCATCATCCCGTATGAAGGTCTGAGCGGCGACCAAATGGGCCAGATCGAAAAAATTTTTAAGGTGGTGTACCCTGTGGATGATCATCACTTTAAGGTGATCCTGCACTATGGCACACTGGTAATCGACGGGGTTACGCCGAACATGATCGACTATTTCGGACGGCCGTATGAAGGCATCGCCGTGTTCGACGGCAAAAAGATCACTGTAACAGGGACCCTGTGGAACGGCAACAAAATTATCGACGAGCGCCTGATCAACCCCGACGGCTCCCTGCTGTTCCGAGTAACCATCAACGGAGTGACCGGCTGGCGGCTGTGCGAACGCATTCTGGCGTAA

**EPG-NanoLuc FFNP** – Vector: pcDNA3.1 – Promoter: CAG

Start-NLS-LgBiT-Linker-EPG-Linker-SmBiT-End

ATGGGCCCTAAAAAGAAGCGTAAAGTCGTCTTCACACTCGAAGATTTCGTTGGGGACTGGGAACAGACAGCCGCCTACAACCTGGACCAAGTCCTTGAACAGGGAGGTGTGTCCAGTTTGCTGCAGAATCTCGCCGTGTCCGTAACTCCGATCCAAAGGATTGTCCGGAGCGGTGAAAATGCCCTGAAGATCGACATCCATGTCATCATCCCGTATGAAGGTCTGAGCGCCGACCAAATGGCCCAGATCGAAGAGGTGTTTAAGGTGGTGTACCCTGTGGATGATCATCACTTTAAGGTGATCCTGCCCTATGGCACACTGGTAATCGACGGGGTTACGCCGAACATGCTGAACTATTTCGGACGGCCGTATGAAGGCATCGCCGTGTTCGACGGCAAAAAGATCACTGTAACAGGGACCCTGTGGAACGGCAACAAAATTATCGACGAGCGCCTGATCACCCCCGACGGCTCCATGCTGTTCCGAGTAACCATCAACGGAGGAGGCGGTAGTCTTACCTGTAACACATGCTCAGTGAGTCTGATTGGAATATGTCTGAATCCCGCAACAGCGACTTGCTCCACCAACACATCCGTCTGCACCACAGGAAGAGCCAGTTTCACGGGCGTCCTCGGCTTCCTGGGCTTCAACTCCCAGGGCTGCACGGAGGGAGCTCAGTGTAATGGCACCGTGTCCGGGTCCATCCTGGGTGCGTCGTACACGGTCACTCAAACCTGCTGCAGCACAAACAACTGCAACCCCGTGACCAGCGGCGCCTCCGGAGGAGGCGGCTCCGTGACCGGCTGGCGGCTGTGCGAACGCATTCTGGCGTAA

**EPG-NanoLuc FR86** – Vector: pcDNA3.1 – Promoter: CAG

Start-NLS-LgBiT-Linker-EPG-Linker-SmBiT-End

ATGGGCCCTAAAAAGAAGCGTAAAGTCGTCTTCACACTCGAAGATTTCGTTGGGGACTGGGAACAGACAGCCGCCTACAACCTGGACCAAGTCCTTGAACAGGGAGGTGTGTCCAGTTTGCTGCAGAATCTCGCCGTGTCCGTAACTCCGATCCAAAGGATTGTCCGGAGCGGTGAAAATGCCCTGAAGATCGACATCCATGTCATCATCCCGTATGAAGGTCTGAGCGCCGACCAAATGGCCCAGATCGAAGAGGTGTTTAAGGTGGTGTACCCTGTGGATGATCATCACTTTAAGGTGATCCTGCCCTATGGCACACTGGTAATCGACGGGGTTACGCCGAACATGCTGAACTATTTCGGACGGCCGTATGAAGGCATCGCCGTGTTCGACGGCAAAAAGATCACTGTAACAGGGACCCTGTGGAACGGCAACAAAATTATCGACGAGCGCCTGATCACCCCCGACGGCTCCATGCTGTTCCGAGTAACCATCAACGGAGGAGGCGGTAGTCTTACCTGTAACACATGCTCAGTGAGTCTGATTGGAATATGTCTGAATCCCGCAACAGCGACTTGCTCCACCAACACATCCGTCTGCACCACAGGAAGAGCCAGTTTCACGGGCGTCCTCGGCTTCCTGGGCTTCAACTCCCAGGGCTGCACGGAGGGAGCTCAGTGTAATGGCACCGTGTCCGGGTCCATCCTGGGTGCGTCGTACACGGTCACTCAAACCTGCTGCAGCACAAACAACTGCAACCCCGTGACCAGCGGCGCCTCCCCTGCCCCAGCTCCCGTGTCCGGCTGGCGGCTGTTCAAGAAAATTTCTTAA

**EPG-NanoLuc RF86** – Vector: pcDNA3.1 – Promoter: CAG

Start-NLS-LgBiT-Linker-EPG-Linker-SmBiT-End

ATGGGCCCTAAAAAGAAGCGTAAAGTCGTCTTCACACTCGAAGATTTCGTTGGGGACTGGGAACAGACAGCCGCCTACAACCTGGACCAAGTCCTTGAACAGGGAGGTGTGTCCAGTTTGCTGCAGAATCTCGCCGTGTCCGTAACTCCGATCCAAAGGATTGTCCGGAGCGGTGAAAATGCCCTGAAGATCGACATCCATGTCATCATCCCGTATGAAGGTCTGAGCGCCGACCAAATGGCCCAGATCGAAGAGGTGTTTAAGGTGGTGTACCCTGTGGATGATCATCACTTTAAGGTGATCCTGCCCTATGGCACACTGGTAATCGACGGGGTTACGCCGAACATGCTGAACTATTTCGGACGGCCGTATGAAGGCATCGCCGTGTTCGACGGCAAAAAGATCACTGTAACAGGGACCCTGTGGAACGGCAACAAAATTATCGACGAGCGCCTGATCACCCCCGACGGCTCCATGCTGTTCCGAGTAACCATCAACCCTGCCCCAGCTCCCCTTACCTGTAACACATGCTCAGTGAGTCTGATTGGAATATGTCTGAATCCCGCAACAGCGACTTGCTCCACCAACACATCCGTCTGCACCACAGGAAGAGCCAGTTTCACGGGCGTCCTCGGCTTCCTGGGCTTCAACTCCCAGGGCTGCACGGAGGGAGCTCAGTGTAATGGCACCGTGTCCGGGTCCATCCTGGGTGCGTCGTACACGGTCACTCAAACCTGCTGCAGCACAAACAACTGCAACCCCGTGACCAGCGGCGCCTCCGGAGGAGGCGGCTCCGTGTCCGGCTGGCGGCTGTTCAAGAAAATTTCTTAA

**EPG-NanoLuc RF114** – Vector: pcDNA3.1 – Promoter: CAG

Start-NLS-LgBiT-Linker-EPG-Linker-SmBiT-End

ATGGGCCCTAAAAAGAAGCGTAAAGTCGTCTTCACACTCGAAGATTTCGTTGGGGACTGGGAACAGACAGCCGCCTACAACCTGGACCAAGTCCTTGAACAGGGAGGTGTGTCCAGTTTGCTGCAGAATCTCGCCGTGTCCGTAACTCCGATCCAAAGGATTGTCCGGAGCGGTGAAAATGCCCTGAAGATCGACATCCATGTCATCATCCCGTATGAAGGTCTGAGCGCCGACCAAATGGCCCAGATCGAAGAGGTGTTTAAGGTGGTGTACCCTGTGGATGATCATCACTTTAAGGTGATCCTGCCCTATGGCACACTGGTAATCGACGGGGTTACGCCGAACATGCTGAACTATTTCGGACGGCCGTATGAAGGCATCGCCGTGTTCGACGGCAAAAAGATCACTGTAACAGGGACCCTGTGGAACGGCAACAAAATTATCGACGAGCGCCTGATCACCCCCGACGGCTCCATGCTGTTCCGAGTAACCATCAACCCTGCCCCAGCTCCCCTTACCTGTAACACATGCTCAGTGAGTCTGATTGGAATATGTCTGAATCCCGCAACAGCGACTTGCTCCACCAACACATCCGTCTGCACCACAGGAAGAGCCAGTTTCACGGGCGTCCTCGGCTTCCTGGGCTTCAACTCCCAGGGCTGCACGGAGGGAGCTCAGTGTAATGGCACCGTGTCCGGGTCCATCCTGGGTGCGTCGTACACGGTCACTCAAACCTGCTGCAGCACAAACAACTGCAACCCCGTGACCAGCGGCGCCTCCGGAGGAGGCGGCTCCGTGACGGGATACAGACTGTTCGAGGAGATCCTTTAA

**EPG-NanoLuc fFFNP** – Vector: pcDNA3.1 – Promoter: CAG

Start-NLS-LgBiT-Linker-EPG-Linker-SmBiT-End

ATGGGCCCTAAAAAGAAGCGTAAAGTCGTCTTCACACTCGAAGATTTCGTTGGGGACTGGGAACAGACAGCCGCCTACAACCTGGACCAAGTCCTTGAACAGGGAGGTGTGTCCAGTTTGCTGCAGAATCTCGCCGTGTCCGTAACTCCGATCCAAAGGATTGTCCGGAGCGGTGAAAATGCCCTGAAGATCGACATCCATGTCATCATCCCGTATGAAGGTCTGAGCGCCGACCAAATGGCCCAGATCGAAGAGGTGTTTAAGGTGGTGTACCCTGTGGATGATCATCACTTTAAGGTGATCCTGCCCTATGGCACACTGGTAATCGACGGGGTTACGCCGAACATGCTGAACTATTTCGGACGGCCGTATGAAGGCATCGCCGTGTTCGACGGCAAAAAGATCACTGTAACAGGGACCCTGTGGAACGGCAACAAAATTATCGACGAGCGCCTGATCACCCCCGACGGCTCCATGCTGTTCCGAGTAACCATCAACGGAGGAGGCGGTAGTAAGTGTGTACTTTTGGGATTCGCAGCAGTGATCGGATTCTTCGCGATCGCGGAGTCTCTTACCTGTAACACATGCTCAGTGAGTCTGATTGGAATATGTCTGAATCCCGCAACAGCGACTTGCTCCACCAACACATCCGTCTGCACCACAGGAAGAGCCAGTTTCACGGGCGTCCTCGGCTTCCTGGGCTTCAACTCCCAGGGCTGCACGGAGGGAGCTCAGTGTAATGGCACCGTGTCCGGGTCCATCCTGGGTGCGTCGTACACGGTCACTCAAACCTGCTGCAGCACAAACAACTGCAACCCCGTGACCAGCGGCGCCTCCTACGTCCAGATCTCCGTCAGCGCGGCCCTGAGCGCCGCCCTGCTGGCCTGCGTCTGGGGCCAGTCCGTCTACGGAGGAGGCGGCTCCGTGACCGGCTGGCGGCTGTGCGAACGCATTCTGGCGTAA

**EPG-NanoLuc fFF86** – Vector: pcDNA3.1 – Promoter: CAG

Start-NLS-LgBiT-Linker-EPG-Linker-SmBiT-End

ATGGGCCCTAAAAAGAAGCGTAAAGTCGTCTTCACACTCGAAGATTTCGTTGGGGACTGGGAACAGACAGCCGCCTACAACCTGGACCAAGTCCTTGAACAGGGAGGTGTGTCCAGTTTGCTGCAGAATCTCGCCGTGTCCGTAACTCCGATCCAAAGGATTGTCCGGAGCGGTGAAAATGCCCTGAAGATCGACATCCATGTCATCATCCCGTATGAAGGTCTGAGCGCCGACCAAATGGCCCAGATCGAAGAGGTGTTTAAGGTGGTGTACCCTGTGGATGATCATCACTTTAAGGTGATCCTGCCCTATGGCACACTGGTAATCGACGGGGTTACGCCGAACATGCTGAACTATTTCGGACGGCCGTATGAAGGCATCGCCGTGTTCGACGGCAAAAAGATCACTGTAACAGGGACCCTGTGGAACGGCAACAAAATTATCGACGAGCGCCTGATCACCCCCGACGGCTCCATGCTGTTCCGAGTAACCATCAACGGAGGAGGCGGTAGTAAGTGTGTACTTTTGGGATTCGCAGCAGTGATCGGATTCTTCGCGATCGCGGAGTCTCTTACCTGTAACACATGCTCAGTGAGTCTGATTGGAATATGTCTGAATCCCGCAACAGCGACTTGCTCCACCAACACATCCGTCTGCACCACAGGAAGAGCCAGTTTCACGGGCGTCCTCGGCTTCCTGGGCTTCAACTCCCAGGGCTGCACGGAGGGAGCTCAGTGTAATGGCACCGTGTCCGGGTCCATCCTGGGTGCGTCGTACACGGTCACTCAAACCTGCTGCAGCACAAACAACTGCAACCCCGTGACCAGCGGCGCCTCCTACGTCCAGATCTCCGTCAGCGCGGCCCTGAGCGCCGCCCTGCTGGCCTGCGTCTGGGGCCAGTCCGTCTACGGAGGAGGCGGCTCCGTGTCCGGCTGGCGGCTGTTCAAGAAAATTTCTTAA

**EPG-NanoLuc fFR86** – Vector: pcDNA3.1 – Promoter: CAG

Start-NLS-LgBiT-Linker-EPG-Linker-SmBiT-End

ATGGGCCCTAAAAAGAAGCGTAAAGTCGTCTTCACACTCGAAGATTTCGTTGGGGACTGGGAACAGACAGCCGCCTACAACCTGGACCAAGTCCTTGAACAGGGAGGTGTGTCCAGTTTGCTGCAGAATCTCGCCGTGTCCGTAACTCCGATCCAAAGGATTGTCCGGAGCGGTGAAAATGCCCTGAAGATCGACATCCATGTCATCATCCCGTATGAAGGTCTGAGCGCCGACCAAATGGCCCAGATCGAAGAGGTGTTTAAGGTGGTGTACCCTGTGGATGATCATCACTTTAAGGTGATCCTGCCCTATGGCACACTGGTAATCGACGGGGTTACGCCGAACATGCTGAACTATTTCGGACGGCCGTATGAAGGCATCGCCGTGTTCGACGGCAAAAAGATCACTGTAACAGGGACCCTGTGGAACGGCAACAAAATTATCGACGAGCGCCTGATCACCCCCGACGGCTCCATGCTGTTCCGAGTAACCATCAACGGAGGAGGCGGTAGTAAGTGTGTACTTTTGGGATTCGCAGCAGTGATCGGATTCTTCGCGATCGCGGAGTCTCTTACCTGTAACACATGCTCAGTGAGTCTGATTGGAATATGTCTGAATCCCGCAACAGCGACTTGCTCCACCAACACATCCGTCTGCACCACAGGAAGAGCCAGTTTCACGGGCGTCCTCGGCTTCCTGGGCTTCAACTCCCAGGGCTGCACGGAGGGAGCTCAGTGTAATGGCACCGTGTCCGGGTCCATCCTGGGTGCGTCGTACACGGTCACTCAAACCTGCTGCAGCACAAACAACTGCAACCCCGTGACCAGCGGCGCCTCCTACGTCCAGATCTCCGTCAGCGCGGCCCTGAGCGCCGCCCTGCTGGCCTGCGTCTGGGGCCAGTCCGTCTACCCTGCCCCAGCTCCCGTGTCCGGCTGGCGGCTGTTCAAGAAAATTTCTTAA

**EPG-NanoLuc fRR114** – Vector: pcDNA3.1 – Promoter: CAG

Start-NLS-LgBiT-Linker-EPG-Linker-SmBiT-End

ATGGGCCCTAAAAAGAAGCGTAAAGTCGTCTTCACACTCGAAGATTTCGTTGGGGACTGGGAACAGACAGCCGCCTACAACCTGGACCAAGTCCTTGAACAGGGAGGTGTGTCCAGTTTGCTGCAGAATCTCGCCGTGTCCGTAACTCCGATCCAAAGGATTGTCCGGAGCGGTGAAAATGCCCTGAAGATCGACATCCATGTCATCATCCCGTATGAAGGTCTGAGCGCCGACCAAATGGCCCAGATCGAAGAGGTGTTTAAGGTGGTGTACCCTGTGGATGATCATCACTTTAAGGTGATCCTGCCCTATGGCACACTGGTAATCGACGGGGTTACGCCGAACATGCTGAACTATTTCGGACGGCCGTATGAAGGCATCGCCGTGTTCGACGGCAAAAAGATCACTGTAACAGGGACCCTGTGGAACGGCAACAAAATTATCGACGAGCGCCTGATCACCCCCGACGGCTCCATGCTGTTCCGAGTAACCATCAACCCTGCCCCAGCTCCCAAGTGTGTACTTTTGGGATTCGCAGCAGTGATCGGATTCTTCGCGATCGCGGAGTCTCTTACCTGTAACACATGCTCAGTGAGTCTGATTGGAATATGTCTGAATCCCGCAACAGCGACTTGCTCCACCAACACATCCGTCTGCACCACAGGAAGAGCCAGTTTCACGGGCGTCCTCGGCTTCCTGGGCTTCAACTCCCAGGGCTGCACGGAGGGAGCTCAGTGTAATGGCACCGTGTCCGGGTCCATCCTGGGTGCGTCGTACACGGTCACTCAAACCTGCTGCAGCACAAACAACTGCAACCCCGTGACCAGCGGCGCCTCCTACGTCCAGATCTCCGTCAGCGCGGCCCTGAGCGCCGCCCTGCTGGCCTGCGTCTGGGGCCAGTCCGTCTACCCTGCCCCAGCTCCCGTGACGGGATACAGACTGTTCGAGGAGATCCTTTAA
